# Visual threats reduce blood-feeding and trigger escape responses in *Aedes aegypti* mosquitoes

**DOI:** 10.1101/2022.01.08.475512

**Authors:** Nicole E. Wynne, Karthikeyan Chandrasegaran, Lauren Fryzlewicz, Clément Vinauger

## Abstract

The diurnal mosquitoes *Aedes aegypti* are vectors of several arboviruses, including dengue, yellow fever, and Zika viruses. To find a host to feed on, they rely on the sophisticated integration of olfactory, visual, thermal, and gustatory cues reluctantly emitted by the hosts. If detected by their target, this latter may display defensive behaviors that mosquitoes need to be able to detect and escape. In humans, a typical response is a swat of the hand, which generates both mechanical and visual perturbations aimed at a mosquito. While the neuro-sensory mechanisms underlying the approach to the host have been the focus of numerous studies, the cues used by mosquitoes to detect and identify a potential threat remain largely understudied. In particular, the role of vision in mediating mosquitoes’ ability to escape defensive hosts has yet to be analyzed. Here, we used programmable visual displays to generate expanding objects sharing characteristics with the visual component of an approaching hand and quantified the behavioral response of female mosquitoes. Results show that *Ae. aegypti* is capable of using visual information to decide whether to feed on an artificial host mimic. Stimulations delivered in a LED flight arena further reveal that landed females *Ae. aegypti* display a stereotypical escape strategy by taking off at an angle that is a function of the distance and direction of stimulus introduction. Altogether, this study demonstrates mosquitoes can use isolated visual cues to detect and avoid a potential threat.

**Summary Statement:** In isolation, visual stimuli programmed to mimic a human swat prevent mosquitoes from blood-feeding by triggering take-offs and escape responses.

## Introduction

Mosquitoes are responsible for transmitting disease-causing pathogens killing more than 700,000 people every year (World Health Organization, 2018). The underlying motivation for adult females of epidemiologically problematic mosquito species is their need for nutrients, including proteins, contained in vertebrate blood to produce progeny. Therefore, the reproductive fitness of female mosquitoes is not only directly linked to their ability to detect, locate and feed on a host but also to survive interactions with a larger and potentially defensive organism. But accessing a resource that is hidden under the skin of mobile and defensive hosts is not free of any risks and, in addition to physiological stresses associated with the ingestion of blood, hosts display defensive behaviors aimed at deterring or killing mosquitoes (Wynne et al., 2020).

Mosquitoes rely on the integration of multiple sensory inputs to find hosts, where olfactory cues gate and modulate responses to visual and thermal targets (Liu and Vosshall, 2019; San Alberto et al., 2021; van Breugel et al., 2015; Vinauger et al., 2019; Zhan et al., 2021). However, how mosquitoes detect defensive maneuvers from their hosts remains largely understudied. In humans, a typical response to biting insects is swatting. From the insect’s perspective, a swat corresponds to a rapidly approaching appendage, most often a hand, on an interception course. When approaching, the appendage induces rapid air displacement (*i*.*e*., mechanical component) and a rapid expansion of an object in the mosquito visual field (*i*.*e*., visual component). Using a mammalian tail simulator, Matherne *et al*., showed that the airflow generated by the swinging of mammals’ tails reduced the proportion of *Aedes aegypti* landing by 50% (Matherne et al., 2018). In free flight, both *Ae. aegypti* and the nocturnal malaria vector *Anopheles coluzzii*, displayed rapid escape maneuvers when stimulated with an artificial swatter that induces airflow and visual cues (Cribellier et al., 2021), hinting at the role of visual cues in signaling threats. However, to our knowledge, no study has yet isolated visual cues from mechanical cues to characterize the role of vision in mosquito escape behaviors.

Utilizing visual cues to detect and assess defensive hosts would allow mosquitoes to categorize potential hosts as defensive from a safe distance or under highly turbulent conditions where air displacement might not be a reliable indicator of a threat. In addition, some species display a “pre-biting” rest (Reid, 1968) or “pre-attack” resting (Clements, 1999) behavior when the mosquito density increases around a target host (Tuno et al., 2003), which is known to increase the amount of host defensive behaviors (Day and Edman, 1984; Edman et al., 1972). With an increase in host defenses but ultimately an endless supply of blood, it would appear adaptive for the mosquito to wait until the host activity calms. During this “pre-biting” phase, among the cues available to mosquitoes to evaluate the host’s defensive behaviors, vision would allow them to do so from a safe distance away and even under highly turbulent conditions where mechanical cues may be unreliable (Tuno et al., 2003; Tuno et al., 2017). In this context, we tested the hypothesis that day-time biting *Ae. aegypti* mosquitoes can use visual cues to detect potential threats and adjust their behavior accordingly. To test our hypothesis, we relied on a quantitative analysis of *Ae. aegypti*’s behavior to investigate how they respond to predator-like looming (expanding) stimuli designed to mimic the visual properties of a slapping hand since humans are *Ae. aegypti’s* preferred host (Christophers, 1960; Ponlawat and Harrington, 2005).

## Materials and Methods

### Mosquitoes

Wild type *Aedes aegypti* mosquitoes (Rockefeller strain, MR-734, MR4, ATCC®, Manassas, VA, USA) were used throughout the experiments. The colony was maintained in a climatic chamber set at 25±1°C, 60±10% relative humidity (RH), and under a 12-12h light-dark cycle. Cages of adults were fed weekly using an artificial feeder (D.E. Lillie Glassblowers, Atlanta, Georgia; 2.5 cm internal diameter) with heparinized bovine blood (Lampire Biological Laboratories, Pipersville, PA, USA) heated at 37°C using a circulating water-bath. Between blood-meals, mosquitoes were fed *ad libitum* with 10% sucrose. Eggs were collected from blood-fed females and hatched in deionized water. Larvae were reared in groups of 200 in covered pans (26×35×4 cm) containing deionized water and fed *ad libitum* with fish food (Hikari Tropic 382 First Bites - Petco, San Diego, CA, USA). Pupae, in groups of 100, were isolated in 16 oz containers (Mosquito Breeder Jar, Bioquip Products, Rancho Dominguez, CA, USA) until emergence.

For all the experiments, 6-8 days old female mosquitoes were used. These females were kept in the presence of males, fed *ad libitum* with 10% sucrose until 24h prior to the experiments, and were never blood-fed. This gave mosquitoes the time to mate in the containers before the experiments (random dissection of randomly selected females revealed that 95% of them had oocytes) and *Ae. aegypti* females of this age class are known to actively seek hosts for blood feeding (Grant and O’Connell, 2007; Tallon et al., 2019). All experiments were performed during the last four hours of the mosquitoes’ subjective day, *i*.*e*., *Zeitgeber Time* (ZT) 8-12 (Eilerts et al., 2018; Taylor and Jones, 1969).

### Expanding stimuli

To present ecologically relevant stimuli, we video-tracked a slapping hand on a collision course with a fictive mosquito located at the camera’s position (GoPro Hero 6 in “linear” recording mode at 60 fps, GoPro, San Mateo, CA, USA). Tracking of the hand’s width at the widest point below knuckle height over ten replicates provided an average duration and expansion velocity profile of a slap (ImageJ; National Institutes of Health, USA; Fig. 1). This information was used to program the symmetrical expansion of a dark square over a bright green background. A square was selected over other shapes as behavioral responses to expanding squares have been shown in tethered *Ae. aegypti females* (Vinauger et al., 2019), and the response characteristics of classical models such as *Drosophila melanogaster* have been extensively analyzed (Tammero and Dickinson, 2002). Furthermore, in the programmable LED arena, a square allowed for the expansion of the stimulus along both dimensions with a higher step resolution than, for example, a disk. The expansion velocity of the fictive swats used throughout the experiment was kept constant at 28.9 horizontal degrees.s^−1^ (LCD monitor) and 183.6 horizontal degrees.s^−1^ (LED arena) as seen from the center of the mosquito container. These virtual swats correspond to the expansion velocity of a hand at 80ms and 40ms before contact, respectively. The rationale behind this choice was to simulate earlier phases of the swat in the feeding assays and later moments (*i*.*e*., closer to impact) in the take-off experiments, as the distance from the display was greater in the feeding assays than in the take-off experiments (300 mm versus 118.5 mm, respectively).

**Figure 1.**
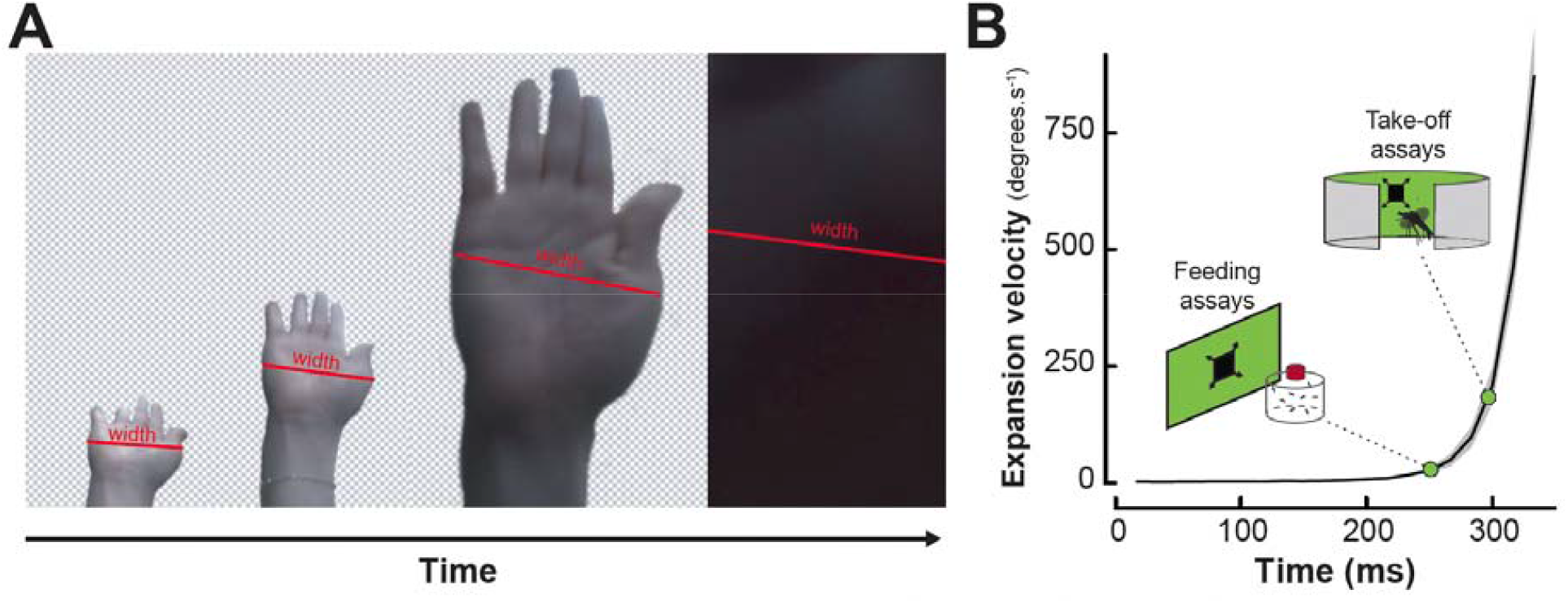
Developing an artificial visual swat. **(A)** Representative screenshots from the video recording of hand slaps filmed at 60 frames per second. The red lines illustrate the measurement of the widest part of the hand. Front view recordings were paired with side-view recordings used to quantify the linear velocity of the hand (data not shown). **(B)** Mean expansion velocity (in degrees per second) of a hand swat as a function of time (black line; n = 10 replicates). Grey shaded area indicates the standard error to the mean expansion velocity. Green points indicate the expansion velocities used to program th expanding squares displayed in the feeding and take-off assays, respectively.

### Artificial blood-feeding assays

#### a) Procedure

To quantify the effect of visual motion on the ability of *Ae. aegypti* females to land and feed on an artificial feeder, we performed blood-feeding assays in the presence of visual patterns generated with PsychoPy v3.0 displayed on a 29-inch (73 cm diagonal) LED-backlit LCD monitor (UltraSharp 29 Ultrawide Monitor - U2917W, Dell, Round Rock, TX, USA). PsychoPy is a Python-based application that allows for the design of behavioral experiments with precise spatial and temporal control of stimuli (Peirce et al., 2019). Specifically, two patterns were generated: 1) a uniform green screen (negative control), and 2) expanding squares (treatment group) programmed to have the same duration as a human swat (Fig. 2A, Movie 1). Green, with a peak irradiance at 535 nm, was chosen as a background color to allow comparisons with the conditions in the LED arena (peak irradiance: 571 nm) (Fig. S1).

**Figure 2.**
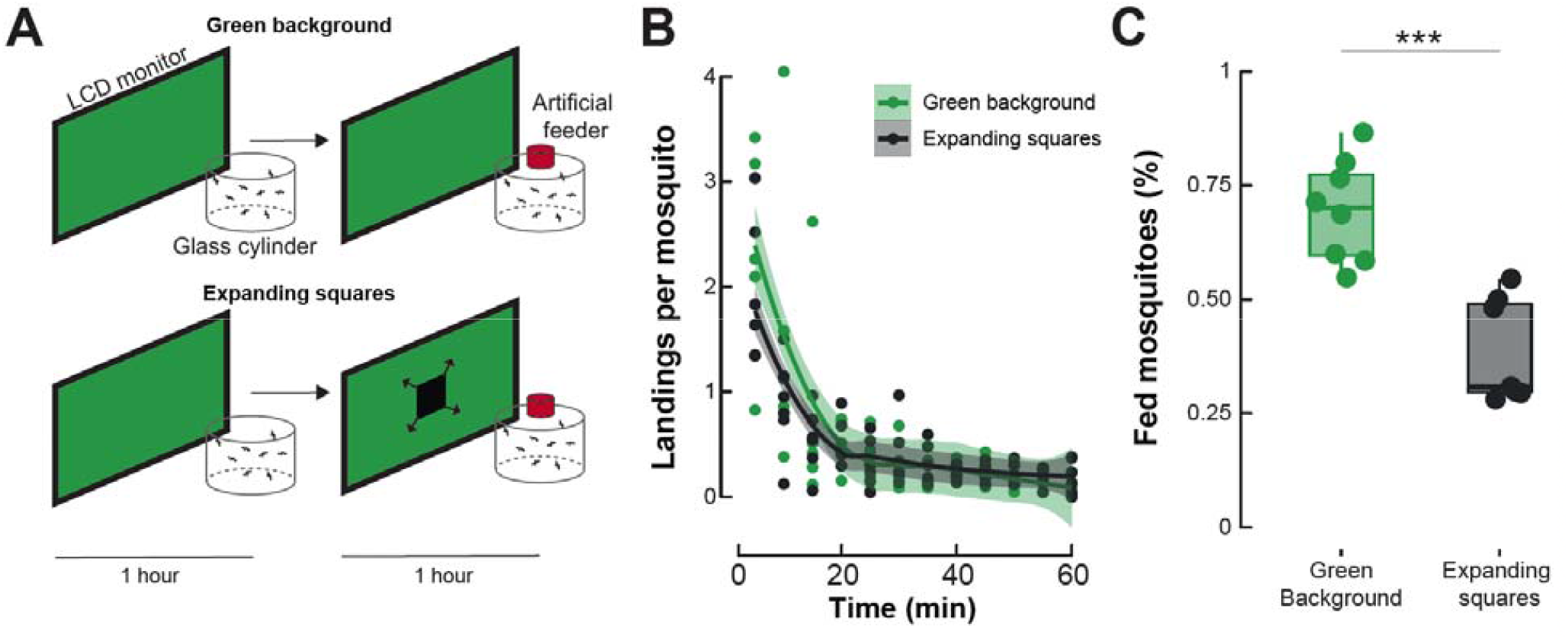
Landing and feeding on a host mimic in different visual contexts. **(A)** Schematic of the experimental apparatus. Groups of mosquitoes are enclosed in a glass cylinder positioned in front of an LCD monitor. To acclimate to the experimental conditions, mosquitoes are left unperturbed in front of a green background for 1 hour. Control groups (top) were exposed to a second hour of green background during which the number of landing and feedings on a membrane feeder were quantified. Treatment groups (bottom) were exposed to randomly positioned squares that symmetrically expand at periodic intervals. **(B)** Number of landings per mosquito quantified every five minutes of the second hour of the experiment for the control (green) and treatment groups (dark grey). **(C)** Proportion of feedings quantified at the end of the second hour of the experiment for the control (green, n = 8) and treatment groups (dark grey, n = 7). Asterisks denote significant differences (***: *p* < 0.001; Generalized linear model).

The mosquitoes used for these experiments (21 < n < 35) were isolated into clear glass cylinders (10 × 10 cm) and starved from sucrose for 24 hours prior to the experiment. Mosquitoes were tested during the last 2 hours of the photophase (ZT 10-12). In the first hour of the experiment, the screen was kept uniformly green to allow mosquitoes to acclimate to the experimental conditions. At the onset of the second hour, a membrane blood feeder was placed on the fabric mesh-lined top side of the glass cylinder. The feeder was warmed to 37□ using a circulating water bath thirty minutes prior to the onset of the experiment. The warmed feeder was filled with ∼ 5 ml of heparinized bovine whole blood (Lampire Biological Laboratories, Pipersville, PA, USA) fifteen minutes prior to the beginning of the experiment to allow the blood to heat up. The fabric mesh-lined side of the clear glass cylinder allowed mosquitoes to sense the heat and see the visual contrast of the feeder. The number of mosquitoes that landed and blood-fed on the feeder were recorded using a camera (Logitech C920, Logitech, Lausanne, Switzerland).

In the negative controls, the baseline levels of landing and feeding were quantified in front of a uniform green screen. In the first hour, mosquitoes were allowed to acclimatize to the experimental conditions including the uniform green background. In the second hour, after placing the blood feeder on a glass cylinder enclosure, the number of landings and feedings on the feeder were quantified. In the treatment groups, mosquitoes isolated in the glass cylinder enclosure were acclimatized to the uniform green screen in the first hour, following which the visual stimuli, *i*.*e*., expanding squares, were introduced every 0.8 seconds for a duration of 0.3 seconds. The positions of the looming squares displayed were randomly determined for each experiment trial to account for potential spatial bias. For this purpose, PsychoPy used an array of cartesian coordinates randomized using the *rand* function in Microsoft Excel.

The glass cylinders were kept in the camera’s field of view throughout the two-hour duration of every experiment trial.

#### b) Statistical analysis

After every trial, the number of engorged females was quantified via visual inspection of their abdomen for the presence of blood, and the number of landings was quantified from the video recordings. These numbers were then compared between treatments as proportions (categorical fixed predictors with two levels: *uniform background* and *looming squares*). For the analysis, we used a Generalized Linear Model assuming a quasibinomial error distribution for the proportion of feeding and a Poisson error distribution for the number of landings per mosquito. The analysis was performed in R (version 3.6.2) using the packages *lme4* (version 1.1-27.1;(Bates et al., 2014)) and *multcomp* (version 1.4-17;(Hothorn et al., 2008)).

### Free-flight LED arena

#### a) Procedure

Experiments were performed in an LED-based arena (*sensu* (Reiser and Dickinson, 2008)) consisting of an array of 96 × 16 LEDs subtending 360° horizontally and 54° vertically from the center of the arena. Individual mosquitoes were cold anesthetized on ice and separated in clear acrylic cylinders with a clear acrylic lid on the top and a fabric mesh lining at the bottom. These containers were kept in the climatic chamber for at least two hours to allow the mosquitoes to recover from the cold anesthesia. Containers were moved to the experimental room (23±1°C, 45±5% RH) thirty minutes before the start of the experiment to allow the mosquitoes to acclimatize to the ambient conditions. In every experimental trial that lasted for thirty minutes, an acrylic cylinder containing a single mosquito was placed inside the arena, with the expanding stimulus being introduced every minute at a randomized angle around the arena (Fig. 3A, Movie 2). The distance and the direction to the point of stimulus introduction were a function of both the mosquito’s position in the arena at the time of introduction and the randomized angle of introduction. The behavior of mosquitoes in the experiment trials was recorded at 30fps with a camera (Logitech C920, Logitech, Lausanne, Switzerland).

**Figure 3.**
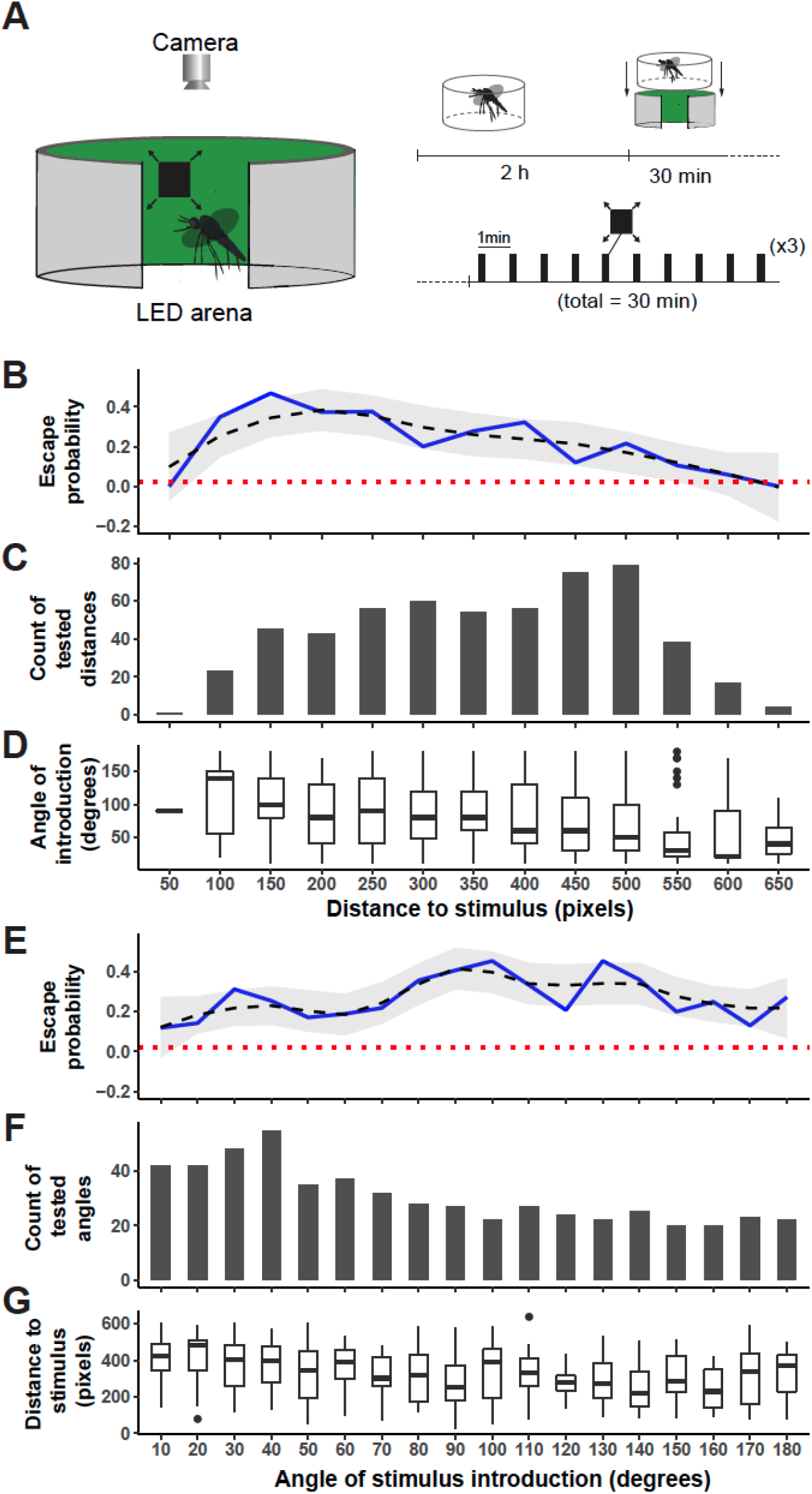
Escape probability as a function of the distance and direction of the stimulus. **(A)** Schematic of the experimental apparatus (left) and graphical representation of th experiment’s timeline (right) where an individual female mosquito is isolated in a clear cylinder 2 hours before the start of the experiments. The cylinder is positioned in the LED arena and the mosquito is provided 30 minutes to acclimatize to the conditions before being stimulated with a looming square every minute for 30 minutes. **(B**,**E)** Mosquitoes’ probability to escape (take-off) after introduction of the expanding stimuli, represented as a function of the distance to the point of stimulus introduction **(B)** or as a function of the direction of stimulus introduction **(E)**. A trendline (dashed, black) from a single-term linear model fit with 95% confidence interval (gray) summarizes the relationship between escape probability and the predictor variables. The dotted horizontal line (red) denotes the baseline escape probability (take-off) in the absence of any expanding stimuli. **(C**,**F)** Number of experimental trials. **(D**,**G)** Distribution of the direction of stimuli introduction (degrees). In **(B-D)**, the response variables are visualized as a function of the mosquitoes’ distance from the point of stimulus introduction (0-650 pixels or 0-177.14 mm) represented as a categorical variable, *i*.*e*., 50 pixels per category. In **(E-G)**, the response variables are visualized as a function of the angle of stimulus introduction (0-180 degrees, where 0 degrees is behind and 180 in front of the mosquito) represented as a categorical variable, *i*.*e*., 10 degrees per category.

#### b) Statistical analysis

Mosquitoes that did not move throughout the duration of the experiment (∼7% of the tested mosquitoes) were determined “not active” and discarded from the analysis. Because of the study’s focus on take-off responses, trials where mosquitoes were in flight before the introduction of the stimulus were discarded from the analysis (n = 63). Based on these criteria, a total of 551 stimuli introductions were conserved, and videos of active landed mosquitoes were trimmed 30 seconds before and 30 seconds after each stimuli introduction. These one-minute trimmed videos were then converted into image sequences and imported into Fiji ImageJ (National Institutes of Health) for manual tracking, using the *Manual Tracking* plugin. The head of each mosquito was tracked before, during, and after stimulus introduction. The stimuli edges and center points were also tracked for each stimulus introduction. From these stimuli coordinates, the mosquito-stimulus distance and the angle of stimulus introduction were recorded. Distances were measured in pixels and the punctual conversions to mm provided throughout the manuscript were obtained by reporting the measured diameter of the arena to its measurement on the videos.

In addition, to calculate the escape probability of mosquitoes we considered responsive mosquitoes as those that took off within 10 frames post stimulus introduction (*i*.*e*., ⅓ of a second), but overall observed that all mosquitoes that responded to the stimulus took off within the duration of the stimulus presentation, and no take offs were observed post stimulus. Take offs past this time frame were not considered responses as mosquitoes would have been intercepted by the fictive object beyond that time.

A baseline probability of take-off was determined by repeating the experiments with the exception that no visual looming stimuli were introduced. All LEDs of the arena were kept ON for thirty minutes and the behavior of individualized mosquitoes (n = 10) was recorded with the same video camera as the treatment group. For each mosquito, ten time points were randomly selected within these 30 minutes, using the sample function in R (*i*.*e*., N = 100). At each of these times and for up to five seconds afterwards, spontaneous take-offs were quantified and used to calculate the baseline take-off probability in the absence of visual stimuli.

All data were saved as .csv and imported in R (version 3.6.2) and trajectories were rotated so that all stimuli introductions were fictively re-positioned at 180°. All the points before the stimulus introduction were considered *pre-stimulus*, while all the points after the stimulus introduction were considered *post-stimulus*. From this data, four response variables were calculated by subtracting the *pre-stimulus* period from the *post-stimulus* period (to account for mosquitoes walking before stimulus introduction): linear and angular velocity, escape direction and displacement: *i*) the linear velocity was determined by the distance between consecutive points divided by the sampling interval (pixels.sec^−1^); *ii*) The angular velocity was calculated as change in heading between consecutive points divided by the sampling interval (degrees.sec^−1^); *iii*) The displacement (distance travelled) and *iv)* escape direction (degrees) were quantified by measuring differences between the mosquito’s location at time of stimulus introduction and its location 20 frames post-stimulus. The effects of the distance and direction to the point of stimulus introduction on these four variables were assessed by means of Generalized Linear Models (GLMs) using the *R* package *lme4* (Bates et al., 2014).

In order to quantify the amount of digitizing error introduced by manual tracking of the mosquitoes, 3 experimenters tracked the same 2-second-long sequence (59 frames), once a day during 3 consecutive days. This provided an estimate of the inter- and intra-individual errors in the digitized cartesian coordinates, quantified here as the Root Square Mean Error (RMSE), calculated according to equation (1):

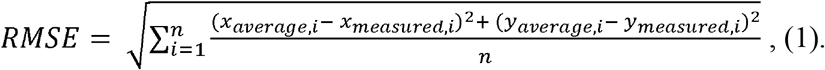

On average, the tracked location of the mosquito deviated by 7.86 pixels (Fig. S2) from the mean of the locations measured by all 3 experimenters. This corresponds to 43.9 % of the body length of an average-sized mosquito landed at the bottom of the arena (17.88 ± 0.47 pixels or 4.87 ± 0.12 mm). In this context, and to minimize the influence of errors introduced during the digitization process, changes in velocity and angular velocity were calculated after smoothing the raw data using a low-pass butterworth filter (*pass*.*filt()*, R package *dplR* 1.7.2 (Bunn, 2008)) with a cut-off frequency of 0.25 (Walker, 1998). The four response variables thus quantified were visualized as a function of the mosquitoes’ distance from the point of stimulus introduction (Fig. 5A-D) and the angle of stimulus introduction (Fig. 5E-H) with a trendline (±95% confidence interval) from a single-term local regression model fit summarizing the relationship (*geom_smooth(method = ‘loess’)*, R package *ggplot2* 3.3.5 (Wickham, 2009)).

**Figure 5.**
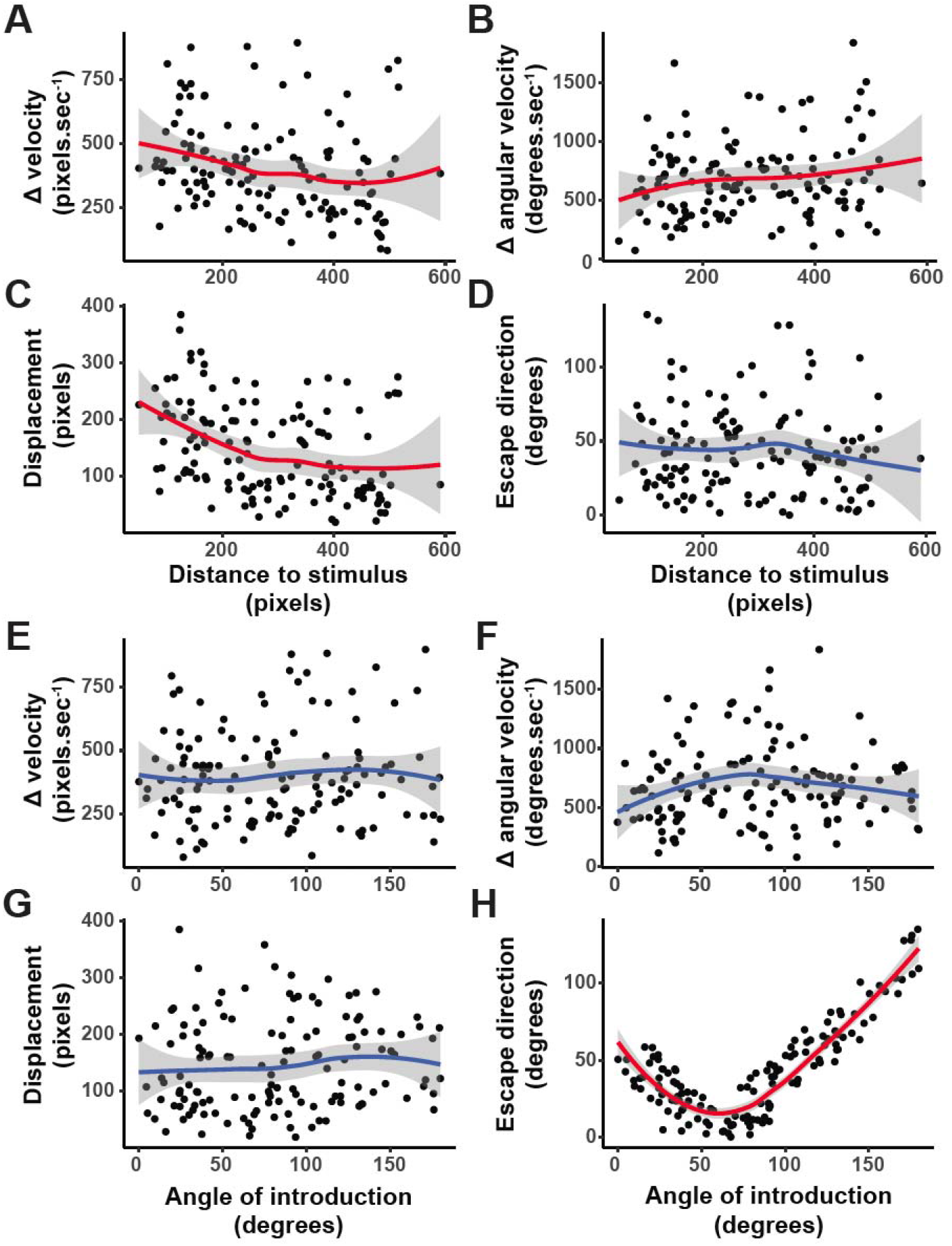
Statistical analysis of the escape strategy displayed by mosquitoes. Mosquitoes’ responses quantified as **(A**,**E)** Change in velocity (pixels.sec^−1^), **(B**,**F)** Change in angular velocity (degrees.sec^−1^), **(C**,**G)** Displacement (pixels), and **(D**,**H)** Direction of escape (degrees), where 180 is directly towards the stimulus and 0 away from the stimulus. In **(A-D)**, the response variables are visualized as a function of the mosquitoes’ distance from the point of stimulus introduction (0-600 pixels or 0-163.5 mm). In **(E-H)**, the response variables are visualized as a function of the angle of stimulus introduction, where 180 is in front of the mosquito and 0 is behind the mosquito. In **(A-H)**, a trend line from a single-term local regression model fit with 95% confidence interval (gray) summarizes the relationship between the response and predictor variables. Red trend lines indicate significant effects of the predictive factors (distance to stimulus or angle of stimulus introduction, respectively) on the response variables, and blue trend lines indicate non-significant effects (Generalized Linear Models, α = 0.05).

## Results

### Extracting visual features of a human swat

Video tracking of a human slapping hand showed an average duration of 335 ± 82 ms with an exponentially increasing expansion rate peaking at 874.9 degrees.sec^−1^ before covering the entire field of view of the camera (n=10; Fig. 1). Side view recordings of 10 slaps, showed that the peak instantaneous velocity of the slap was of 8.01 ± 0.66 m.s^−1^, which is slower, but within the same order of magnitude as the recorded speed of Olympic boxers’ punches (*i*.*e*., ∼9.14 ± 2 m.sec^−1^; (Walilko et al., 2005)). The duration of the swat was used to program a dark looming square displayed to the mosquitoes on either a LCD monitor or on a programmable LED arena (Fig. 2A and 3A, respectively).

### Visual motion reduces feeding, but not landing, on a warm source of blood

To analyze whether visual, threat-like stimuli could impact mosquitoes’ ability to blood-feed, groups of adult females were provided access to a host mimic positioned in front of a visual display. In presence of a uniform green background, a total of 1226 landings on the feeder were observed over the course of the experiment. When looming squares were repeatedly presented to the mosquitoes, a total of 1042 landings were observed over the course of the experiment and, although there was a tendency for mosquitoes to land more during the early phase of the uniform background experiment, the difference between the two experimental conditions was not significant (Generalized linear model: *p* = 0.51; Fig. 2B).

Concerning blood-feedings, on average, 69.6 ± 2.16 % of mosquitoes fed to repletion in front of the green background (Fig. 2C). However, this proportion was significantly reduced when expanding squares were introduced at regular intervals (38.7 ± 3.19 %; Generalized linear model: *p* < 0.001; Fig. 2C).

### Mosquitoes’ response to visual threats is a function of the distance and direction of the stimulus

To better understand the role of looming visual objects in the context of threat avoidance, mosquitoes were stimulated at regular intervals, with the expanding stimulus being introduced at random positions around the arena. Because mosquitoes were free to fly (or not), this randomization led to a large number of randomized directions of stimulus introduction relative to the mosquito, and random distances between the location of the mosquito and the point of stimulus introduction on the arena (Fig. 3B-G). Mosquitoes’ responses to expanding stimuli introduced at all the directions were sampled homogeneously with stimulations right behind the mosquito (<40 degrees) being sampled relatively more than the rest (Supplementary table 1). Likewise, the mosquitoes’ responses as a function of their distance to the expanding stimuli was sampled homogeneously except for positions where the mosquitoes were either too close (<100 pixels or 27.2 mm in distance) to or too far (>550 pixels or 149.8 mm in distance) from the stimuli (Supplementary table 2).

In the absence of any stimulation, the probability of take-offs was 0.02 (Fig. 3B,E). This number significantly increased to 0.26 when an expanding stimulus was introduced (including stimuli delivered from all directions and distances; Binomial Exact test: *p* < 0.001). However, mosquitoes’ escape probability varied significantly as a function of their distance to the stimulus, with peaks for stimuli delivered between 150-200 pixels away from the mosquito (40.8 - 54.5 mm; escape probability = 0.47) (Generalized linear model, *p* < 0.001). While the effect of the angle at which the expanding square was introduced was not significant (Generalized Linear Model, *p* = 0.0817), the probability of escape tented to peak at angles comprised between 80° and 130°, *i*.*e*., laterally to the mosquito.

To confirm that mosquitoes were indeed evading the expanding square, we compared the spatial occupancy of the arena between mosquitoes that were in flight but not stimulated and mosquitoes that took off following stimulus introduction. Because of the randomization of the stimulus introduction, data were rotated to align all stimuli introductions at 180°. While non-stimulated mosquitoes avoided the center of the arena, they explored the entire periphery of the device (Kuiper’s one sample test of uniformity: n.s.; test statistic = 1.4514, critical value at *α* = 0.05: 1.747; Fig. 4A). On the other hand, stimulated mosquitoes not only avoided the center of the arena, but also flew to the sectors directly opposed from where the stimulus was introduced (Kuiper’s one sample test of uniformity: p < 0.05; test statistic = 4.4521 critical value at *α* = 0.05: 1.747; Fig. 4B). To further characterize the escape response displayed by mosquitoes, we analyzed how far mosquitoes would go from the point of stimulus introduction (displacement) and their overall escape direction as a function of both the distance and angle of stimulus introduction. Although there was a tendency for mosquitoes to go further away when the stimulus was introduced closer to them (blue vectors in Fig. 4C), the distance to the point of stimulus introduction had no significant effect on the displacement after take-off, regardless of the angle at which the stimulus was introduced (Fig. 4C). However, the angle of stimulus introduction had a significant effect on the direction of escape (Generalized Linear Model; *p* < 0.001). When the stimulus was introduced in front of the mosquitoes (closer to 180°, red points), they predominantly escaped by performing a slight turn relative to the direction of the stimulus. Mosquitoes that had stimuli introduced laterally (around 90-135°; yellow-orange points) escaped with a wider angle (>90° relative to the origin of the stimulus). Finally, mosquitoes that had stimuli introduced from behind (<90°, blue points) escaped by orienting the most away from the origin of the stimuli (Fig. 4D).

**Figure 4.**
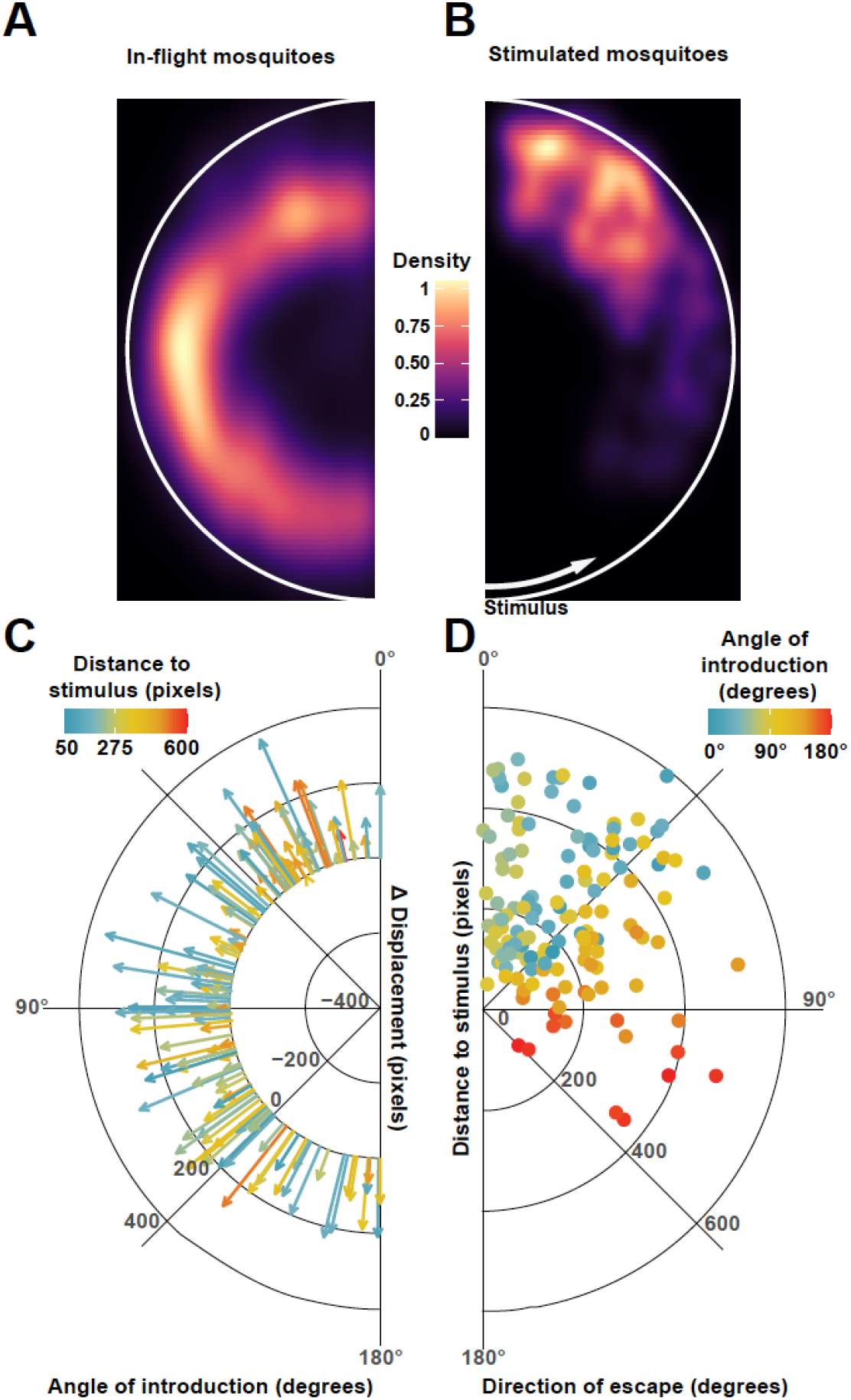
Overall behavioral responses to the looming square. **(A**,**B)** 2-D histogram of the locations occupied by mosquitoes within the LED arena. Data are mirrored on the vertical axis; the white semi circles indicate the relative position of the arena’ walls. **(A)** Normalized histogram of the location of mosquitoes during randomly selected flight bouts, without visual stimulation. **(B)** Normalized histogram of the location of mosquitoes during the first second after a take-off triggered by the introduction of a looming square. Data are oriented such as the stimulus introduction (represented as a white arrow) is positioned at th bottom of the figure. **(C)** Circle plot showing the displacement of mosquitoes (i.e., distance travelled on a scale ranging from 0-400 pixels or 0-109 mm) within 2/3^rd^ of a second of stimulus-triggered take-offs. The length of the vectors encodes for displacement, color coded as a function of the distance to the stimulus’ point of introduction, and the position of the vector around the circle encodes for the direction of stimulus introduction, where 180° is to the front of the mosquito and 0 behind the mosquito. Each vector represents one take-off **(D)** Circle plot representing the direction of escape (position around the circle, where 180 is towards and 0 away from the stimulus) as a function of the distance to the stimulus point of introduction (position along the radius of the circle) and color coded as a function of the direction of stimulus introduction (where 180 is to the front of the mosquito and 0 behind the mosquito). Each point represents one take-off (n = 142).

Analyzing four aspects of the escape response (*i*.*e*., velocity, angular velocity, displacement, and direction of escape) we found that all but the direction of escape were significantly influenced by the distance to the point of stimulus introduction (Generalized Linear Models: *p* = 0.003, *p* = 0.032, *p* < 0.001, and *p* = 0.739 for the velocity, angular velocity, displacement, and direction of escape, respectively. Fig. 5A-D). When analyzed as a function of the angle of stimulus introduction, only the direction of escape was significantly affected (Generalized Linear Models: *p* = 0.82, *p* = 0.31, *p* = 0.59 and *p* < 0.001 for the velocity, angular velocity, displacement, and direction of escape, respectively. Fig. 5E-H). In Fig. 5H, an angle of 180° indicates the front of the mosquito and 0° indicates behind the mosquito, and escape directions of 180° indicate flying towards the stimulus and 0° away from the stimulus. These results show that mosquitoes escaped by flying the most away from the looming stimulus when it was introduced between 45° and 90° or from the back side of the mosquito. When the stimulus was introduced in front of the mosquitoes, they flew towards it but at a slight angle. Remarkably, take-offs were observed even for stimulus introductions immediately behind the mosquito (0-45° angles), highlighting mosquitoes’ capacity to detect threats coming from all directions along the azimuth.

## Discussion

Our results demonstrate that mosquitoes can use isolated visual cues to detect and respond to potential threats, *i*.*e*., even in the absence of other sensory information (*e*.*g*., air displacement due to the host’s movements). In our feeding assays, we found that the presence of looming visual cues did not impact the number of landings on the artificial feeder. However, the proportion of mosquitoes that fed to repletion was reduced by close to 50%. This indicates that while the presence of rapidly moving objects in the visual field of flying mosquitoes does not impair their ability to navigate towards a source of food, once landed these same visual objects trigger an escape response. Although the identity of each mosquito wasn’t maintained in our analysis, the large number of landings relative to the number of feeding events suggests that several landing attempts were made by the same mosquitoes and that several mosquitoes landed (several times) but did not feed.

Using an LED-based flight arena, we were able to further characterize this response by introducing expanding squares at randomized distances and directions relative to the focal mosquito. Results confirmed that looming squares trigger mosquitoes to take-off at a higher probability than chance (*i*.*e*., compared to the take-off probability of landed mosquitoes randomly sampled in the absence of visual stimulation: 0.02). The peak escape probability (0.47) observed here was comparable to the frequency of successful escapes from a mechanical swatter by in-flight *Ae. aegypti* females (Cribellier, 2021). The closer to the mosquito the stimulus was introduced, the more likely it was to take-off and although all directions of introduction induced take-offs, threats originating laterally tended to elicit more escapes. In particular, mosquitoes were most responsive to looming stimuli that came from approximately 30°, 90°, and 130°. This raises the question of whether the heterogeneity of the mosquito retina (Hu et al., 2014) underlies spatial specialization for the detection of threats and warrants further studies to determine the range of elevations above the plane of the substrate at which mosquitoes can detect threats. This is particularly relevant as, here, only mosquitoes that were landed horizontally on the substrate were tested. The field of view available to a mosquito actively engaged in blood-feeding may differ from its resting position (*e*.*g*., different head angle, substrate orientation). In tethered *Drosophila melanogaster*, the expansion of centrally versus laterally positioned objects elicited different response profiles as well, where frontal objects induced strong leg and wing-beat frequency responses but minimal changes in wing-beat amplitudes while lateral objects elicited stronger changes in wing-beat amplitude and transient increases in wing-beat frequency, but did not evoke leg responses (Tammero and Dickinson, 2002; Tammero et al., 2004).

Remarkably, the escape responses we observed were not random: after taking-off, mosquitoes rapidly maneuvered to position themselves out of the line of interception with the trajectory of the fictive threat. Analysis of the trajectories displayed after take-off shows that the direction of escape is indeed directly influenced by the angle of stimulus introduction. A similar behavior had been observed in flight in *D. melanogaster* where looming targets triggered visually-directed banked turns that reoriented the fly’s path within a handful of wingbeats (Muijres et al., 2014). Given the high wing-beat frequency of mosquitoes (400-500Hz), high-speed recordings will be required to quantify the fine scale adjustments performed by mosquitoes to take-off and reorient within such a short timeframe. Additionally, the closer the mosquito was from the point of introduction of the threat, the faster and further they escaped by following a less tortuous path (indicated by a lower angular velocity). These results open new avenues for investigating distance perception by mosquitoes.

It is possible that the response probability to the looming stimuli was actually higher than what we observed by only quantifying take-offs as it may have taken other forms, such as a freeze response to threats. Indeed, other animals have been shown to make a decision to either freeze or flee when threat is presented (Yilmaz and Meister, 2013). Our analysis focused on fleeing responses, given that freezing responses could not be identified and characterized because we focused on landed mosquitoes. In addition, because of the size of the clear acrylic cage the mosquitoes were contained in, escape responses could only be examined over a short period of time, as mosquitoes would rapidly reach the wall of the container and redirect their flight trajectory. However, in spite of this limitation and while quantifying responses in 2-dimensions only, results of the present study show clear visual responses to isolated visual stimuli, warranting further 3-dimensional analysis in larger scale (*e*.*g*., wind tunnel) assays.

The most striking feature of the escape response quantified here is the tight relationship between the angle at which the stimulus is introduced and the direction of escape. Within 2/3rd of a second, mosquitoes did not just take off and scatter away from the stimulus. Conversely, they displayed a clear response pattern that can be preliminarily divided into three categories: 1) when the visual threats appear towards the front of the mosquito, they take off and veer at angle between 45 and 90° away from the origin of the threat; 2) when visual threats appear laterally, mosquitoes take off and re-orient the most away from the threat; 3) when visual threats appear from behind the mosquitoes, they take off and turn slightly, as needed in order to get away from the line of interception.

Altogether, results from the present study provide some insights into the “pre-biting’ or “pre-attack” resting behavior observed in several anopheline and culicine mosquito species (Tuno et al., 2003). Because this “pre-biting” rest was more frequently observed at higher mosquito densities, we had emitted the hypothesis that mosquitoes possessed the ability to evaluate the level of defensiveness from afar, possibly using vision in this context. In this hypothetical scenario, mosquitoes would land near the host and wait for its defensive behaviors to decrease before approaching, thus increasing the probability of a successful blood-feeding. However, in our artificial feeding assays, the number of landings on a host mimic was not affected by visual threats, only the proportion of successful feedings on an otherwise defenseless feeder. This differs from other studies conducted in tabanid flies, where stripes present on the fur of the host made it much more difficult for the flies to land on them (Caro et al., 2019). As a result, horses with solid color experienced more fly landings.

In nature, another critical contributor to mosquitoes’ feeding success is the density of female mosquitoes around the host (Day and Edman, 1984; Edman et al., 1972; Edman et al., 1974; Walker and Edman, 1985). Indeed, as the density of mosquitoes increases so does the level of defensiveness of the host, which negatively correlates with the proportion of females that successfully blood-feed (Edman et al., 1972). As a consequence, mosquitoes prefer to feed on the least defensive or least active host over more defensive ones (Edman et al., 1974; Warnes and Finlayson, 1987). While our results suggest that *Ae. aegypti* gauges the defensiveness of a host only after landing on it, it is worth highlighting that only visual cues were available in our experiments. But while mechanical cues (*i*.*e*., air displacement) generated by a mammal tail simulator significantly prevented mosquitoes from landing (Matherne et al., 2018), whether they can trigger the pre-biting resting behavior remains to be determined. In a previous study, when a mechanical vibration calibrated to mimic an average swat was repeatedly paired with host olfactory cues, *Ae. aegypti* females learned the association between the mechanical stimulus and the odor, and subsequently avoided the trained odor (Vinauger et al., 2018). Mechanical cues thus contribute to host selection processes (Vinauger et al., 2018; Wolff and Riffell, 2018; Wynne et al., 2020) and, by disturbing the ability of mosquitoes to land on the host, most likely contribute to a mosquito’s evaluation of the host’s defensiveness.

Beyond their role when presented in isolation, sensory cues from multiple modalities (*e*.*g*., vision and olfaction) are integrated by the mosquito brain (San Alberto et al., 2021; van Breugel et al., 2015; Vinauger et al., 2019). In the context of evading host defenses, Cribellier *et al*. (Cribellier, 2021) showed that, in flight, *Ae. aegypti* females were most successful at evading a mechanical swatter in bright light conditions, *i*.*e*., when both visual and mechanical cues were available. Interestingly, night-active *Anopheles coluzzii* were most successful at escaping in the dark, suggesting that nocturnal mosquitoes maximize their escape performance by adjusting parameters of their flight behavior itself, such as adopting a more stochastic (*i*.*e*., protean) flight (Cribellier et al., 2021). In the present study, we provide an experimental paradigm that permits the presentation of visual cues in isolation from mechanical perturbations. In future work, this paradigm could be adapted to spatially decouple visual and mechanosensory cues to deepen our understanding of multimodal sensory integration processes in the context of threat avoidance.

Vision has been shown to be used by many insect species in a variety of biological contexts (Olberg, 2012; Palavalli-Nettimi and Theobald, 2020; Palavalli-Nettimi et al., 2019; Spaethe et al., 2006), including identifying predators or threats. Across the animal kingdom, looming stimuli are typically characterized as a threat and elicit an escape response (Peek and Card, 2016). Here, we showed that isolated visual looming stimuli are sufficient to elicit an escape response in *Ae. aegypti* mosquitoes. This highlights the importance of this historically overlooked sensory modality as, in addition to recent studies that characterized its role in the host-seeking context (Liu and Vosshall, 2019; San Alberto et al., 2021; van Breugel et al., 2015; Vinauger et al., 2019; Zhan et al., 2021), vision also contributes to the identification and avoidance of potential threats. The looming stimuli used in the present study were designed to mimic a human swat, as reflecting the preferred host of *Ae. aegypti*. However, mosquito species differ in their host preference, with some species biting smaller or larger mammals (Matherne et al., 2018), amphibians, or birds (Molaei et al., 2008), which vary drastically in their antiparasitic and defensive behaviors. As a consequence, it is likely that the tuning for certain ranges of object sizes, shapes, expansion rates, as well as linear and angular velocities will differ between mosquito species. In *D. melanogaster*, such feature-dependent effects have been described in tethered preparations where taller stripes (above 50° in height) elicited fixation by the animal but shorter stripes (8 to 37° tall) triggered anti-fixation behaviors (Maimon et al., 2008). Compared to flies, mosquitoes face an additional challenge as females are required to approach hosts (larger and slowly expanding visual objects) that can turn into threats at any point (*i*.*e*., swat appendages at high velocities). Without a blood meal, *Ae. aegypti* mosquitoes cannot produce progeny (Christophers, 1960). Given the strong selective pressures associated with host seeking and blood feeding, one might thus expect a similarly fine tuning of the response to features of visual stimuli. Future work investigating this area will provide invaluable insights into the adaptations underlying each species-specific mosquito-host association.

To conclude, results from the present study show that visual threat-like-stimuli alone are sufficient to disrupt *Ae. aegypti*’s blood feeding behavior and trigger a stereotypical escape response influenced by the distance and direction of the threat. This work opens new research avenues for improving our understanding of the role of vision in mosquito biology and developing control strategies that target this sensory modality.

## Acknowledgements

We thank Drs. Chloé Lahondère, Jake Tu, and Jake Socha for their feedback and valuable insights on the experimental design and the manuscript. We also thank Danny Eanes and Steve Lowe for their help with technical details, Chanel Hsu for her help with preliminary data management and visualization, and Shajaesza Diggs and Darren Dougharty for their help with mosquito rearing.

## Author contributions

Conceptualization: N.W., K.C, C.V.; Methodology: N.W., K.C., C.V.; Software: K.C., C.V.; Validation: N.W.; Formal analysis: K.C., C.V.; Investigation: N.W., L.F.; Resources: C.V.; Data Curation: N.W., K.C.; Writing – original draft preparation: N.W., K.C.; Writing – review and editing: N.W., K.C., C.V.; Visualization: N.W., K.C., C.V.; Supervision: C.V.; Project administration: N.W., C.V.; Funding acquisition: C.V.

## Data Availability

Data will be made available from the Dryad Digital Repository (DOI will be inserted here and reference will be added to the list of reference) upon manuscript acceptance.

## Supplementary Information

**Figure S1.**
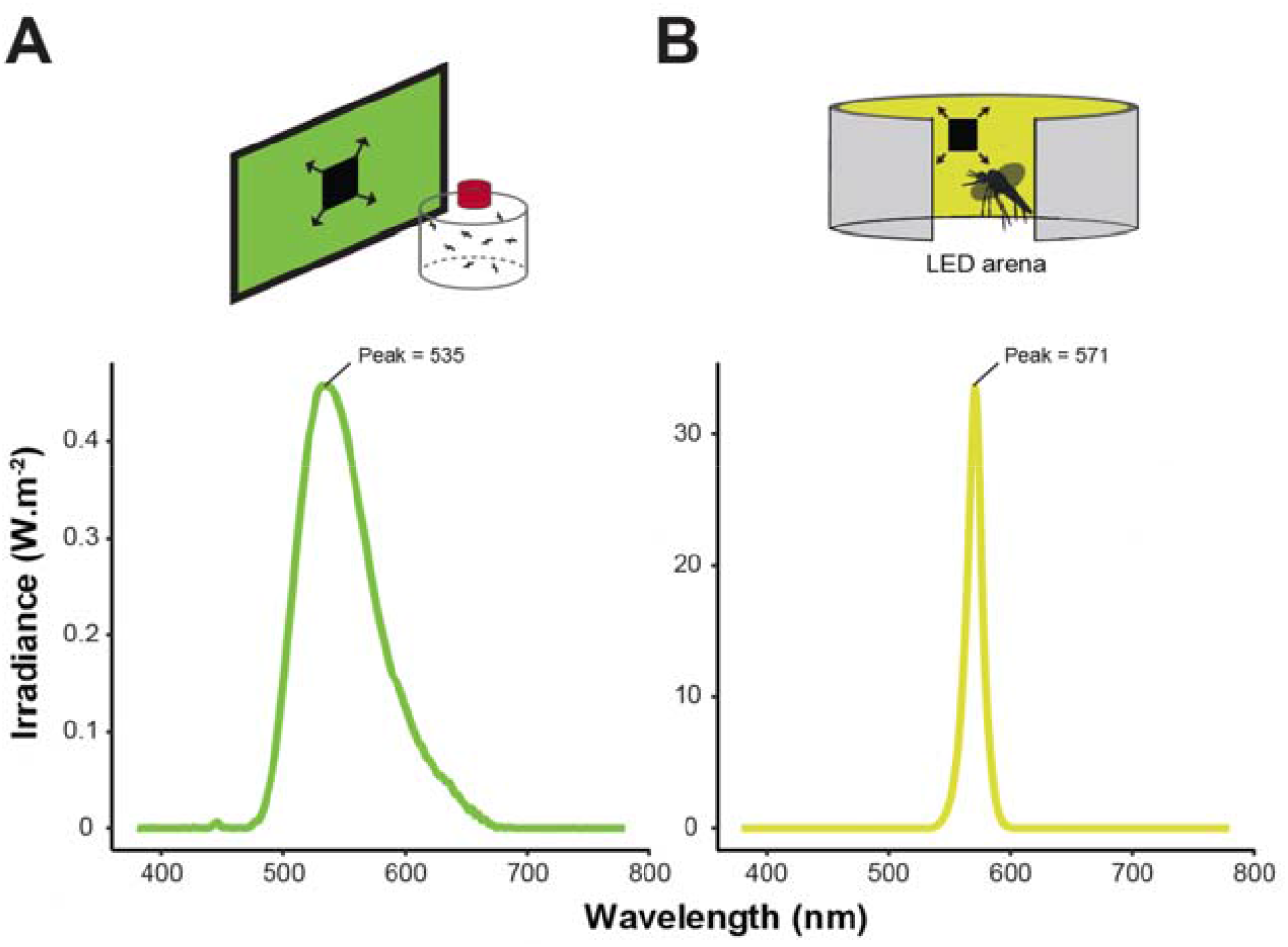
Spectral irradiance of the visual displays. The spectral signature of the green backgrounds used in the feeding **(A)** and take-off **(B)** assays was determined by measuring the irradiance of displays for wavelengths between 380 and 780 nm, using a portable spectral irradiance colorimeter (OHSP350, Hopoocolor, Hangzhou China).

**Figure S2.**
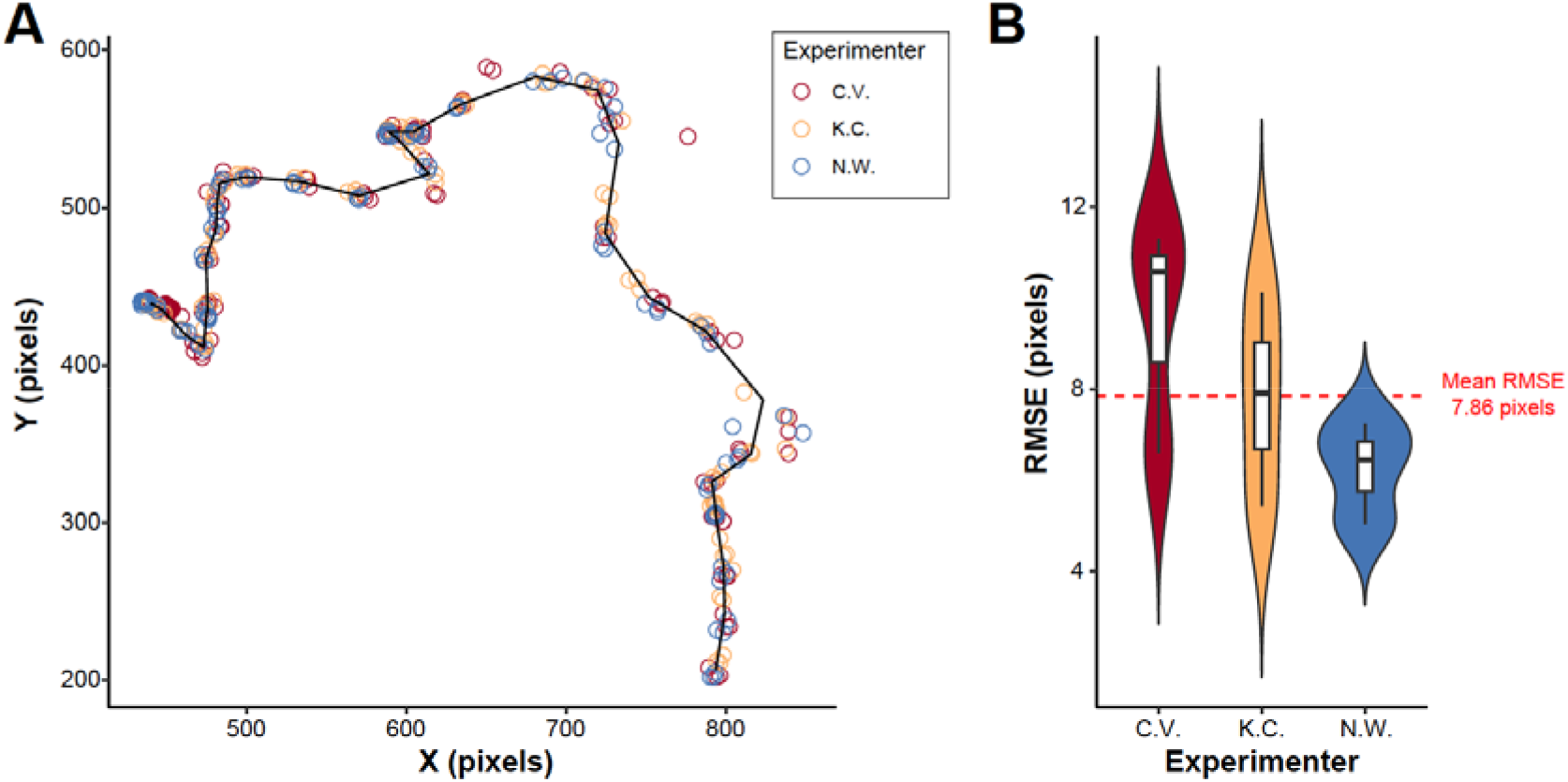
Digitization error associated with manual tracking. **(A)** Tracking of a 59-frame flight sequence by 3 different experimenters on 3 consecutive days. Experimenters are identified by their initials. Data were collected in triplicates and results are color-coded for the individual experimenter. The solid black line indicates the mean trajectory calculated from the mean of the coordinates tracked by all 3 experimenters. **(B)** Digitizing error quantified as the Root Square Mean Error (RMSE), representing the average deviation from the mean trajectory, for each experimenter. The dashed red line indicates the mean RMSE.

**Figure S3.**
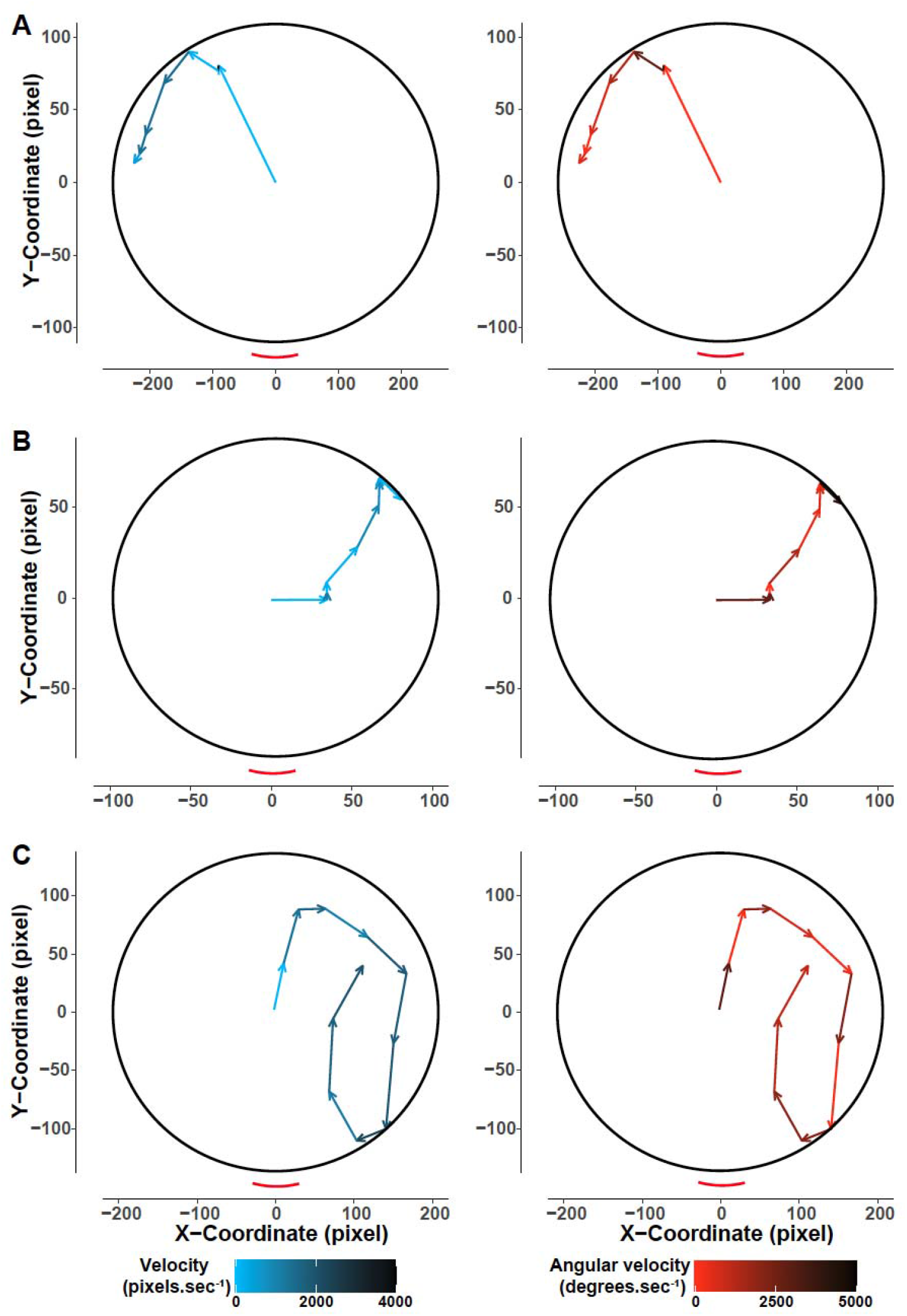
Representative trajectories of mosquitoes’ escape (take-off) responses. Digitized trajectories of the escape (take-off) responses of three mosquitoes **(A-C)** to the expanding stimulus. The visualized cartesian coordinates are tracks of the respective mosquitoes’ head 20 frames post stimulus introduction (*i*.*e*., ⅔ of a second). The trajectories are rotated to fictively re-position the stimuli introductions at 180°. The trajectories are color coded as a function of the mosquitoes’ velocity (left panel, blue) and angular velocity (right panel, red).

**Movie 1. Illustration of stimuli delivered in the two experiments**.

**Movie 2. Example of a mosquito taking-off in response to the looming square**.

**Supplementary Table 1:** Number of mosquitoes sampled for their responses to expanding stimuli as a function of the angle of stimulus introduction.

| **Stimulus direction (degrees)** | **No. trials** | **Lower CI** | **Upper CI** | **z-ratio** |
| --- | --- | --- | --- | --- |
| 0 – 10 | 42 | 31 | 56.8 | 24.223 |
| 10 – 20 | 42 | 31 | 56.8 | 24.223 |
| 20 – 30 | 48 | 36.2 | 63.7 | 26.82 |
| 30 – 40 | 55 | 42.2 | 71.6 | 29.719 |
| 40 – 50 | 35 | 25.1 | 48.7 | 21.034 |
| 50 – 60 | 37 | 26.8 | 51.1 | 21.964 |
| 60 – 70 | 32 | 22.6 | 45.3 | 19.605 |
| 70 – 80 | 28 | 19.3 | 40.6 | 17.632 |
| 80 – 90 | 27 | 18.5 | 39.4 | 17.126 |
| 90 – 100 | 22 | 14.5 | 33.4 | 14.498 |
| 100 – 110 | 27 | 18.5 | 39.4 | 17.126 |
| 110 – 120 | 24 | 16.1 | 35.8 | 15.569 |
| 210 – 130 | 22 | 14.5 | 33.4 | 14.498 |
| 130 – 140 | 25 | 16.9 | 37 | 16.094 |
| 140 – 150 | 20 | 12.9 | 31 | 13.397 |
| 150 – 160 | 20 | 12.9 | 31 | 13.397 |
| 160 – 170 | 23 | 15.3 | 34.6 | 15.037 |
| 170 – 180 | 22 | 14.5 | 33.4 | 14.498 |

**Supplementary Table 2:**
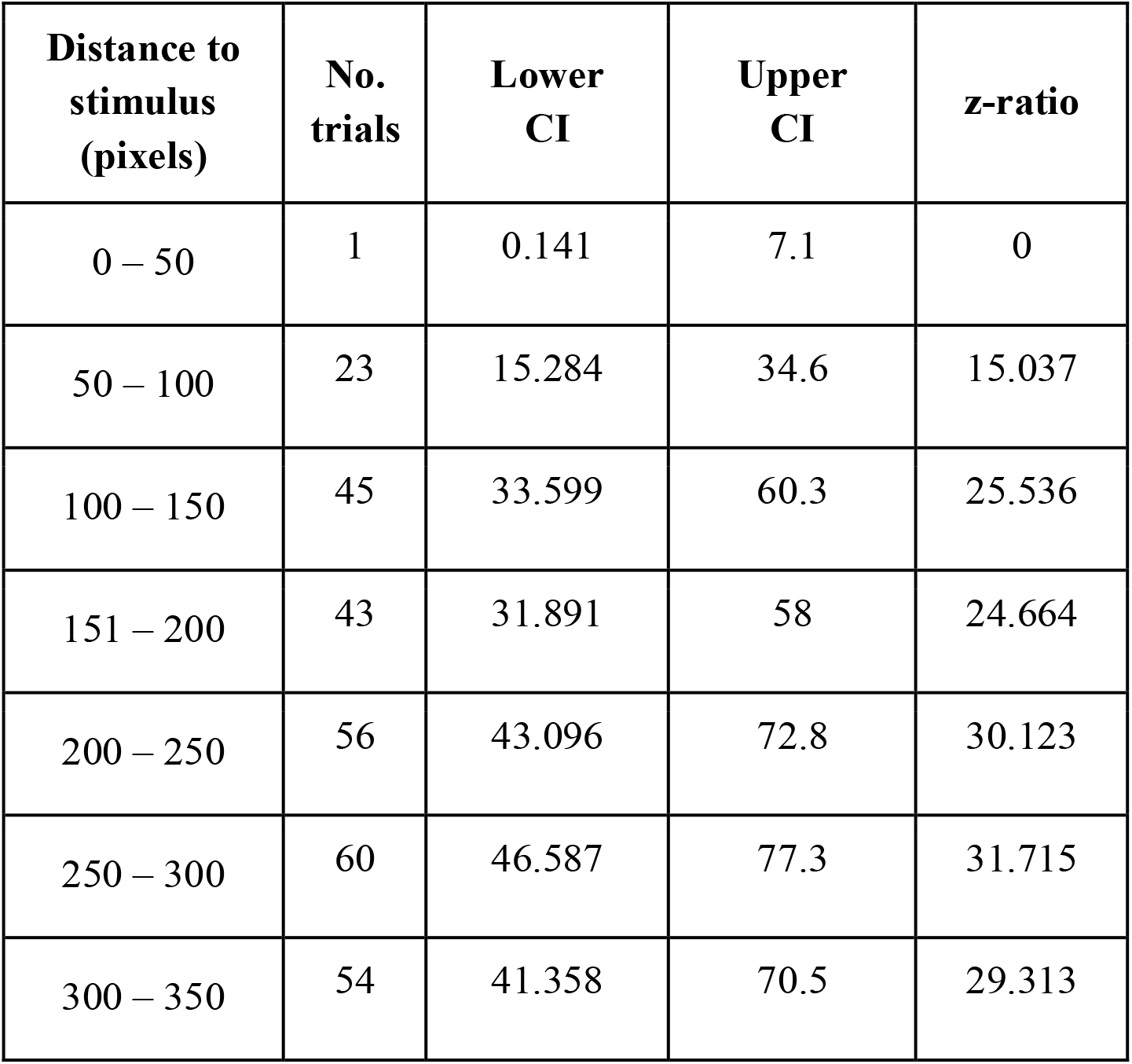
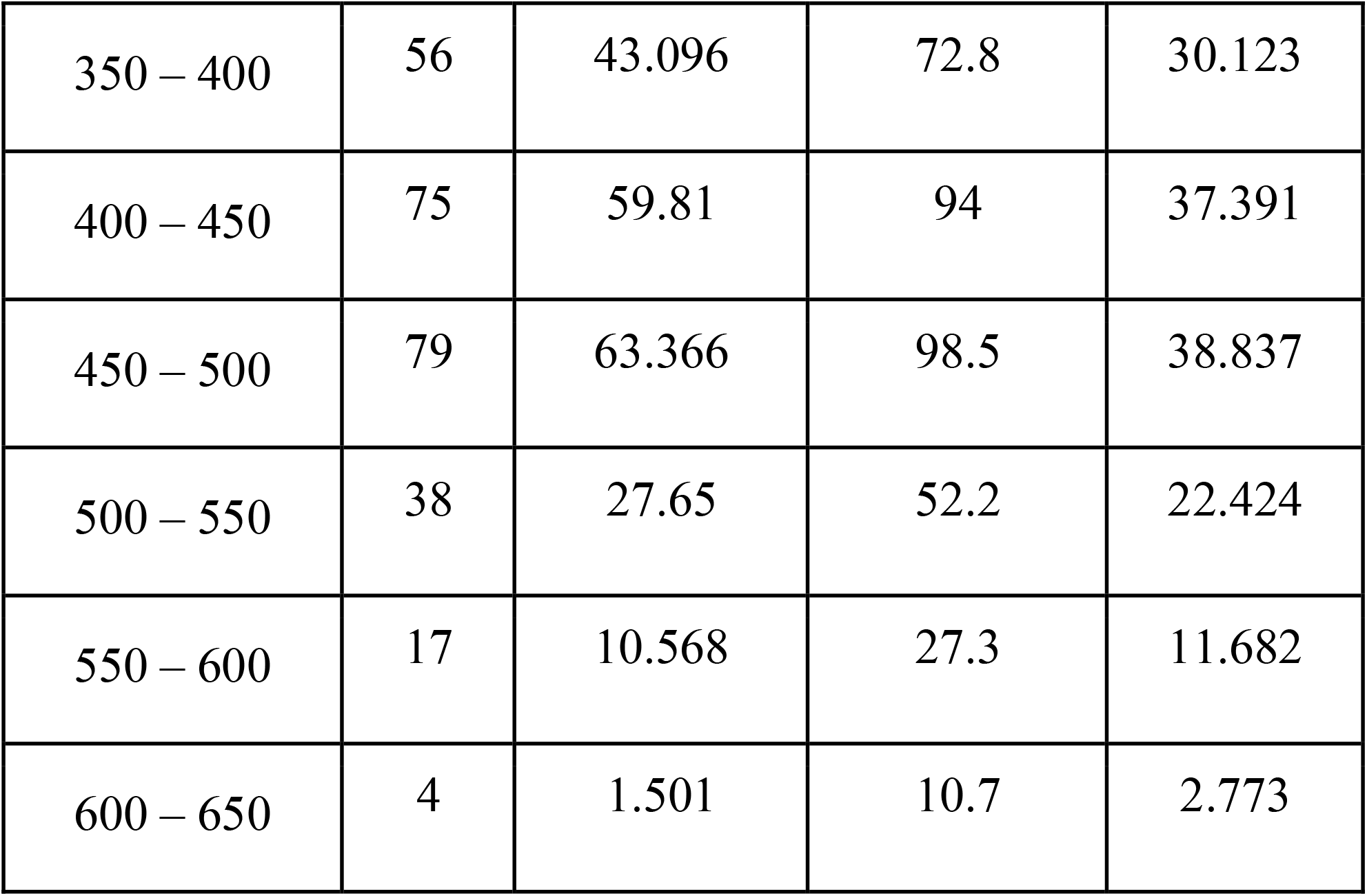
Number of mosquitoes sampled for their responses to expanding stimuli as a function of the mosquitoes’ distance to the expanding stimulus

| **Distance to stimulus (pixels)** | **No. trials** | **Lower CI** | **Upper CI** | **z-ratio** |
| --- | --- | --- | --- | --- |
| 0 – 50 | 1 | 0.141 | 7.1 | 0 |
| 50 – 100 | 23 | 15.284 | 34.6 | 15.037 |
| 100 – 150 | 45 | 33.599 | 60.3 | 25.536 |
| 151 – 200 | 43 | 31.891 | 58 | 24.664 |
| 200 – 250 | 56 | 43.096 | 72.8 | 30.123 |
| 250 – 300 | 60 | 46.587 | 77.3 | 31.715 |
| 300 – 350 | 54 | 41.358 | 70.5 | 29.313 |
| 350 – 400 | 56 | 43.096 | 72.8 | 30.123 |
| 400 – 450 | 75 | 59.81 | 94 | 37.391 |
| 450 – 500 | 79 | 63.366 | 98.5 | 38.837 |
| 500 – 550 | 38 | 27.65 | 52.2 | 22.424 |
| 550 – 600 | 17 | 10.568 | 27.3 | 11.682 |
| 600 – 650 | 4 | 1.501 | 10.7 | 2.773 |

